# The Formin Inhibitor, SMIFH2, Inhibits Members of the Myosin Superfamily

**DOI:** 10.1101/2020.08.30.274613

**Authors:** Yukako Nishimura, Shidong Shi, Fang Zhang, Rong Liu, Yasuharu Takagi, Alexander D. Bershadsky, Virgile Viasnoff, James R. Sellers

## Abstract

The small molecular inhibitor of formin FH2 domains, SMIFH2, is widely used in cell biological studies. It was selected in a chemical screen as a compound inhibiting formin-driven actin polymerization *in vitro*, but not polymerization of pure actin, and found to be active against several types of formins from different species (Rizvi et al., 2009). Here, in experiments with cultured fibroblasts, we found that SMIFH2 inhibits retrograde flow of myosin 2 filaments and contraction of stress fibers. We further checked the effect of SMIFH2 on non-muscle myosin 2A and skeletal muscle myosin 2 *in vitro* and found that SMIFH2 inhibits myosin ATPase activity and ability to translocate actin filaments in the *in vitro* motility assay. While inhibition of myosin 2A *in vitro* required somewhat higher concentration of SMIFH2 than inhibition of retrograde flow and stress fiber contraction in cells, inhibition of several other non-muscle myosin types, e.g. mammalian myosin 10, *Drosophila* myosin 7a and *Drosophila* myosin 5 by SMIFH2, was equally or more efficient than inhibition of formins. Since actin polymerization and myosin contractility are linked in many cytoskeleton processes, additional careful analysis is needed in each case when function of formins was proposed solely on the basis of experiment with SMIFH2.

## Introduction

Formins are a large and diverse class of actin associated proteins that are evolutionarily conserved in nature (Breitsprecher and Goode, 2013; Schönichen and Geyer, 2010; van Gisbergen and Bezanilla, 2013). *In vitro*, formin activities include nucleation and processive elongation of actin filaments (Courtemanche, 2018; Paul and Pollard, 2009; Zigmond, 2004); some formins also bundle actin filaments (Harris et al., 2006; Michelot et al., 2006; Schönichen et al., 2013) and bind to microtubules (Bartolini et al., 2008; Chesarone et al., 2010; Gaillard et al., 2011). Formins contain two types of characteristic domains, formin homology (FH) domains 1 and 2. The FH1 domain contains proline-rich motifs that interact with profilin-actin complex thereby recruiting actin monomers (Courtemanche and Pollard, 2012; Paul et al., 2008). The FH2 domains form dimers, which can nucleate actin filaments and function as processive caps at the filament plus (barbed) ends (Aydin et al., 2018; Courtemanche, 2018; Goode and Eck, 2007; Paul and Pollard, 2009). Combined action of FH1 and FH2 domains strongly accelerates filament growth.

Formins are thought to be required for the many tasks including the formation of filopodia, stress fibers, lamellipodia and cytokinetic rings (Breitsprecher and Goode, 2013; Chhabra and Higgs, 2007; Schönichen and Geyer, 2010). However, because of multiplicity of formins (mammals have 15 genes encoding FH1 and FH2 domains) and apparent redundancy between them, it is often not easy to prove that particular cellular functions depend on formins based on the knockout/knockdown experiments. In addition, in some cases, a rapid inhibition of formin function is necessary. Therefore, a broad specificity chemical formin inhibitor (Rizvi et al., 2009) was widely used in studies of formin functions *in vivo*.

Rizvi et al (2009) conducted a small molecule screen to identify compounds that inhibited the assembly of actin filaments stimulated by the mouse formins, mDia1 and mDia2 in the presence of profilin *in vitro*. A compound termed SMIFH2 was identified that inhibited such assembly in a concentration dependent manner. Half-maximal inhibition of mDia1 occurred at ∼15 µM SMIFH2 concentration. SMIFH2 did not affect assembly of pure actin. At saturating SMIFH2 concentrations the rate of actin assembly equaled that of actin in the absence of formin (Rizvi et al., 2009). Truncation studies suggested that the target of the drug was the FH2 domain. Formins from a variety of species including *C. elegans* CYK-1, *S. pombe* Cdc12, *S. pombe* Fus1, *S. cerevisae* Bni1, and *M. musculus* mDia2 were also inhibited with IC_50_ values ranging from 5-15 µM SMIFH2 suggesting that the inhibitor would be generally applicable to all formins (Rizvi et al., 2009), which however, was not directly checked.

Other inhibitors affecting actin polymerization such as marine toxins latrunculin A and B (Spector et al., 1983), jasplakinolide (Bubb et al., 1994), swinholide A (Bubb et al., 1995), fungal toxins cytochalasins (Natori, 1986), or Amanita mushroom toxin phalloidin (Wieland and Faulstich, 1978) are natural products selected by evolution. High specificity of some of them, e.g. latrunculin A was confirmed in genetic experiments showing that yeasts with mutated actin lacking latrunculin A binding ability are viable at very high concentrations of this drug (Ayscough, 1998; Morton et al., 2000). Others nevertheless can have dual functions, as cytochalasin B, which affects both actin polymerization and glucose transport (Kapoor et al., 2016; MacLean-Fletcher and Pollard, 1980; Yamada and Wessells, 1973). By contrast, the chemical structure of SMIFH2 suggests that this compound can hardly be specific – due to highly electrophilic nature (Baell, 2010) – even though the molecular targets other than formin FH2 domain have not been clearly identified. Of note, off-target effects have been reported *in vivo* such as the alteration of the function of the tumor suppressor protein, p53, albeit at relatively high concentration (Isogai et al., 2015).

Nevertheless, the common belief was that at least in the area related to the cytoskeleton and cell motility, this inhibitor can be safely used for the identification of formin functions. This compound has thus been broadly utilized by the cytoskeleton community to study the role of formin-dependent actin polymerization in a variety of species including human, mouse, chicken, zebrafish, Drosophila, Arabidopsis and yeasts, and in diverse cell types including platelets, fibroblasts, epithelial cells, oocytes, as well as various cancer cells (Isogai et al., 2015).

Our present study shows that SMIFH2 appears to be also a potent inhibitor of molecular motors of myosin family. In the course of cell motility and shape changes, the processes of actin polymerization obviously function in concert with numerous processes mediated by diverse myosin motors. Thus, conclusions about formin involvement in particular cell functions made solely on the basis of experiments with SMIFH2 should be carefully analyzed and perhaps reconsidered.

## Results

### SMIFH2 inhibits contraction of actomyosin fibers and myosin filament flow in living and permeabilized cells

The initial observation which triggered this study was inhibition of traction forces exerted by REF52 fibroblast upon treatement with 30 μM of SMIFH2. The effect was apparent already within 10 minutes following the SMIFH2 addition, when integrity of stress fiber system was still well preserved (Fig.1A). The degree of inhibition of traction forces by SMIFH2 was comparable with that by myosin 2 ATPase inhibitor, para-amino blebbistatin (80 μM) (Fig. 1B).

**Figure 1.**
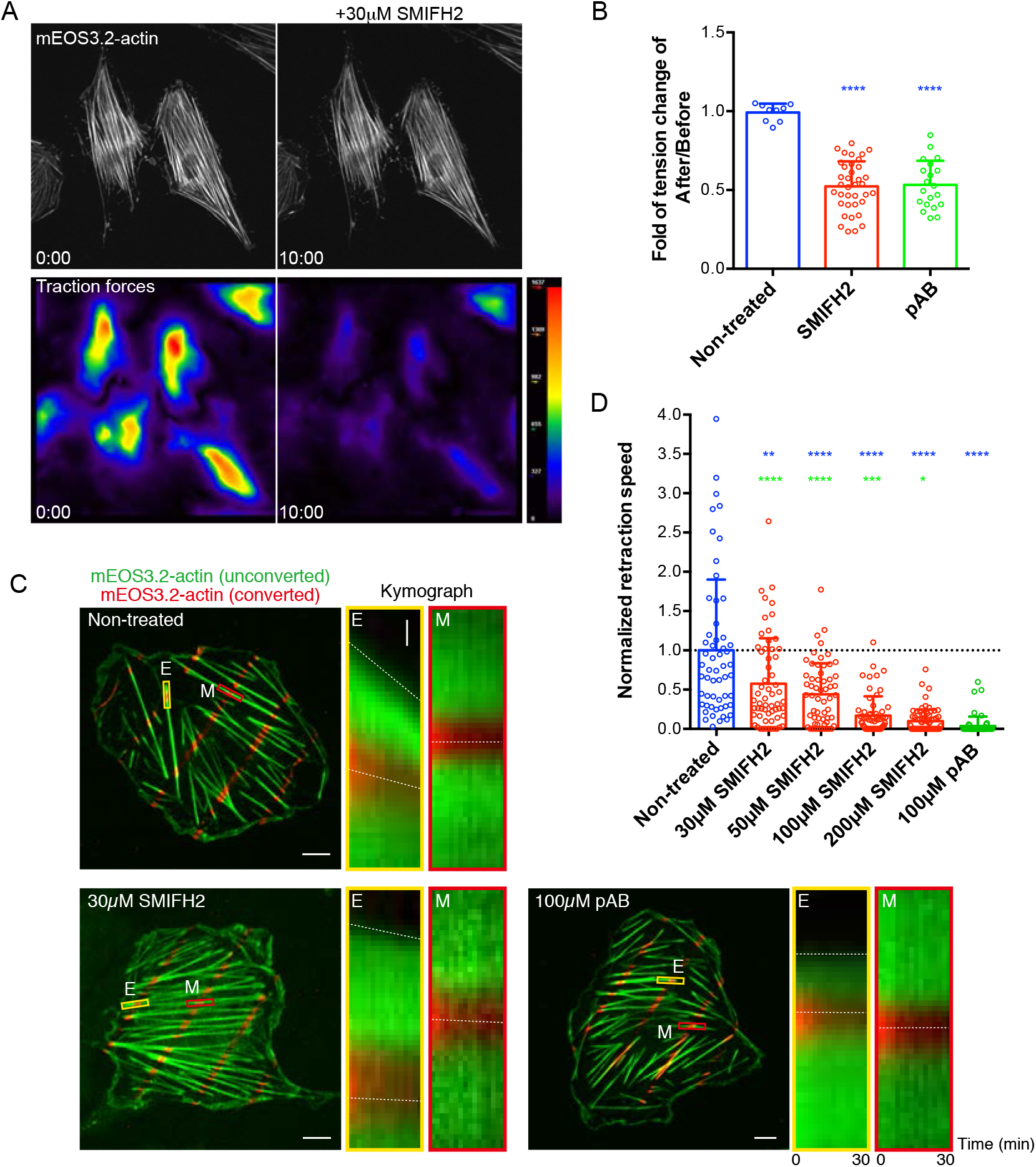
SMIFH2 inhibits stress fiber contractility in living and permeabilized REF52 cells. (A) Reduction of traction forces in living REF52 cells by treatment with SMIFH2. Stress fibers visualized by expression of mEos3.2-actin (top panels) and magnitude of traction forces exerted by cells on substrate shown as heat maps (bottom panels) in cells immediately after (left panels) and 10 minutes following (right panels) addition of 30μM SMIFH2. Note that while overall actin organization did not change, the traction forces dropped dramatically. (B) The quantification of the drop in traction forces upon treatment with 30μM SMIFH2 and 100μM photo-insensitive blebbistatin (pAB). The ratios between values of mean traction forces magnitude per cell at 10 minutes following addition of the drugs to that in the same cells before treatment are presented. The P-values between control group and respective drug treatment groups were calculated using an unpaired two-tailed student t-test. (C) ATP-dependent ventral stress fiber retraction in permeabilized REF52 cells. Cells were labeled by expression of photoconvertible mEOS3.2-actin. Unconverted mEOS3.2-actin is shown in green and pattern of photoconverted mEOS3.2-actin obtained by local laser-illumination is shown in red. Kymographs showing the dynamics of total and photoconverted actin taken at the ends (E, yellow rectangles) or in the middle segments (M, red rectangles) of the ventral stress fibers are shown for each experimental condition. Dashed lines in kymographs demonstrate the movements of the ventral stress fiber ends or photoconverted actin spots. Scale bar, 10 *µ*m. Vertical scale bars in enlarged images of boxed areas, 1 *µ*m. (D) Quantification of the retraction speed of stress fiber ends normalized to the mean speed of non-treated cells (0.025 μm/min). Each dot represents the normalized mean retraction speed of stress fiber ends in one cell. About 40 ends per cell were measured in 56-72 cells under each experimental condition. Bars represent mean ± s.d. The P values were calculated using two-tailed unpaired Student’s t-test. Blue stars indicate the P values for the differences between SMIFH2 and blebbistatin (pAB) treated cell groups and non-treated cell group. Green stars indicate the P values for the differences between SMIFH2 treated and blebbistatin (pAB) treated cell groups. 4, 3, 2, 1 asterisk symbols denote P values P < 0.0001, P < 0.001, P < 0.01, and P < 0.05, respectively. The exact P values are shown in Supplementary Table 1.

We further investigated how SMIFH2 would affect the ATP-dependent contractility of linear ventral stress fibers in REF52 cells. To study the effect of SMIFH2 on actomyosin contraction we used REF52 permeabilized by Triton-X100 (Tee and Bershadsky, 2016; Tee et al., 2015; Tint et al., 1991). Detergent treatment leads to the depletion of all soluble factors from the cells and in particular ATP. Supplementing the medium with ATP induces the myosin 2 dependent contraction of the stress fibers. To monitor the local contraction of the ventral stress fibers at their ends and in the central zone, we expressed photoconvertible mEOS3.2-actin, whose emission wavelength can be converted from green to red channel upon blue laser illumination (Zhang et al., 2012a). We locally photoconverted spots along the length of ventral stress fibers and tracked both the retraction of the unconverted stress fiber ends and the longitudinal movements of the photoconverted actin spots after adding ATP into the solution (Fig. 1C, left panel). We found that ATP addition induced retraction of the stress fibers and centripetal displacement of photoconverted actin spots adjacent to their ends. The retraction speed of the stress fibers tips was, however, faster than the rate of displacement of nearby photoconverted actin spots (Fig. 1C, middle panel), while actin spots in the central zone of stress fibers were hardly mobile (Fig. 1C, right panel). Quantification of the retraction speed of the stress fibers ends revealed that SMIFH2 treatment inhibited their ATP-induced retraction in a dose-dependent manner (Fig. 1C and D). For concentrations of SMIFH2 more than 100 μM (3 fold higher than the typical concentration used in experiments with cells in Rizvi et al (2009), we find the same level of complete inhibition of contractility, comparable to a treatment with para-amino blebbistatin (Fig.1D). At concentrations around 50μM the inhibition is reduced by half but is still significant as compare to control (Fig. 1D).

We tested the inhibition of contraction of another type of contractile acto-myosin structures, the transverse arcs formed by periodically arranged myosin and actin filaments in fibroblasts (Hu et al., 2017). We measured the velocity of movement of transverse arc in human foreskin fibroblast (HFF) cells plated on circular fibronectin island (Tee et al., 2015). In control living cells, non-muscle myosin II filaments visualized by expression of GFP-myosin light chain localized to the transverse arcs and moved toward the cell center with an average velocity of 0.152 μm/min as determined by particle image velocimetry (PIV) (Fig.2A, cf. Hu et al (2017)). Here, we showed that the velocity of this movement decreased in cells treated with SMIFH2 for 45 minutes in dose-dependent manner (Fig.2B; Fig. 2G). Such treatment with SMIFH2 however affected the overall organization of actin and myosin II filaments in the cells (Fig. 2B) in agreement with previous publications (Rizvi et al., 2009; Tee et al., 2015). Permeabilization of the same cells by Triton-X100 removed G-actin and ATP. The centripetal movement of transverse arcs can be induced in permeabilized cells by addition of ATP to the solution (Tee and Bershadsky, 2016; Tee et al., 2015). The treatment with SMIFH2 inhibited the ATP-induced centripetal movement of myosin II filaments in permeabilized cells, even at lowest dose (Fig.2C, D and H). 50 μM of SMIFH2 blocked the movement as efficiently as treatment with para-amino blebbistatin (Fig.2E, F and H).

**Figure 2.**
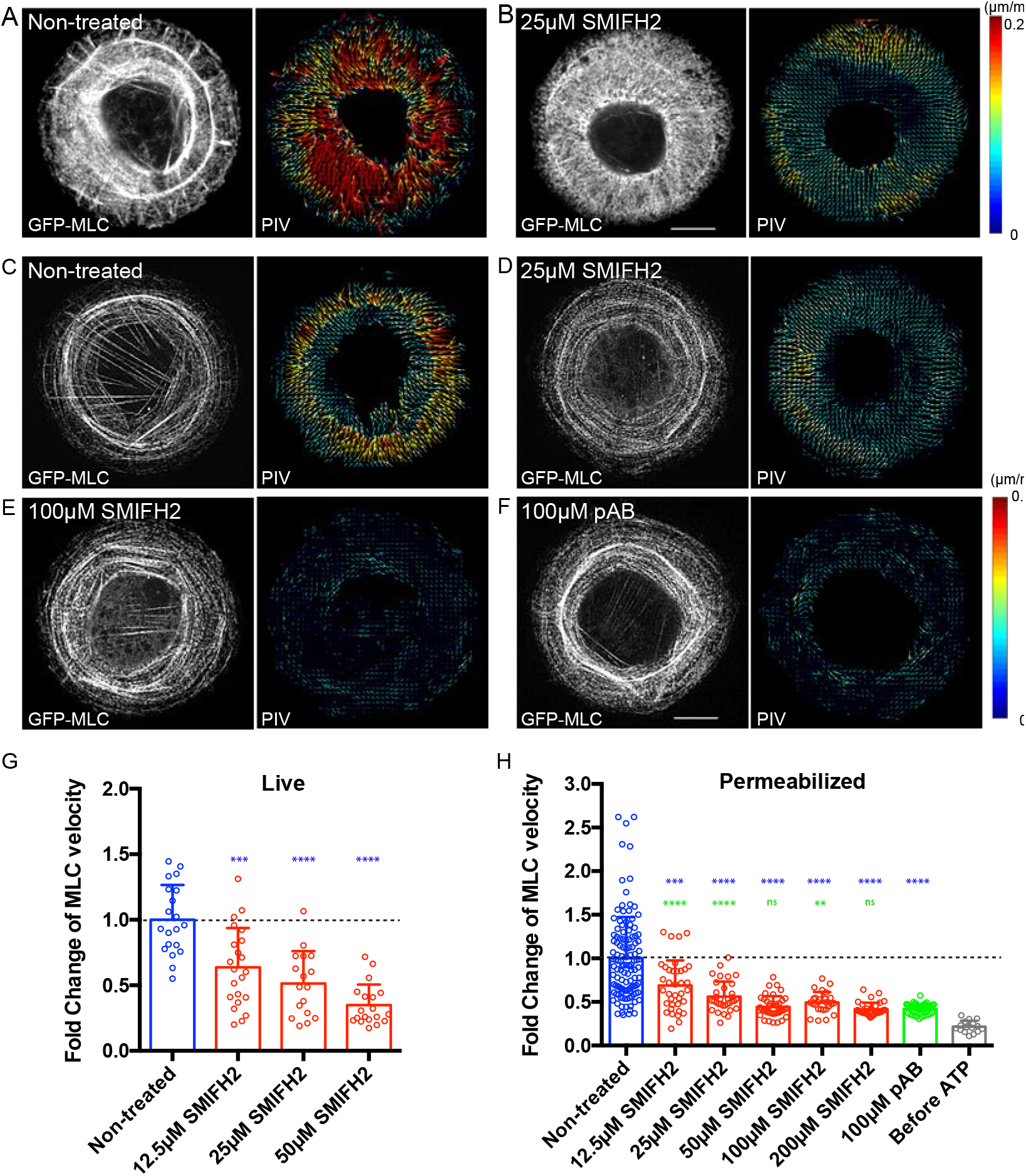
SMIFH2 inhibits centripetal movement of myosin II filaments in living and permeabilized HFF cells. (A, B) Treatment with SMIFH2 reduces the velocity of centripetal movement of myosin II filaments at transvers arc in the live HFF cells plated on circular fibronectin islands. Representative images are shown in non-treated (A) and 25μM SMIFH2-treated cells (B). Myosin II mini-filaments were visualized by expression of GFP-MLC (left panels) and their dynamics were shown as vector maps using particle image velocimetry analysis (PIV, right panels). Arrows represents direction and velocity with color code shown in the right. Scale bar, 10 *µ*m. (C-F) Effects of SMIFH2 and photo-insensitive blebbistatin (pAB) on the velocity of centripetal movement of myosin II filaments induced by ATP in permeabilized HFF cells. Representative images of myosin II filaments (GFP-MLC, left) and their dynamics (PIV, right) are shown in non-treated (C), 25μM (D) or 100μM (E) SMIFH2 treated, and 100μM para-aminoblebbistatin (pAB) treated cells (F). Scale bar, 10 *µ*m. (G) Quantification of the velocity of myosin II filament in non-treated and 12.5, 25 or 50 μM of SMIFH2-treated living cells. Treatment with SMIFH2 reduces the centripetal movement of myosin II filament in a dose-dependent manner. Bars represent mean ±s.d. and each dot represents the value of PIV per cell (n≥17 cells). Values were normalized to the mean speed in non-treated cells (0.152 μm/min). (H) Quantification of the ATP-dependent velocity of myosin II filaments in permeabilized cells with or without pharmacological perturbation. Bars represent mean ±s.d. and each dot represents the mean value of PIV per cell (n≥16 cells). Values were normalized by the mean velocity in non-treated cells (0.0745 μm/min). The P-values calculated using a two-tailed unpaired Student *t*-test are indicated. Blue stars indicate the P values between SMIFH2 or blebbistatin (pAB) treated cells and cells from non-treated group. Green stars indicate the P values between SMIFH2 treated and blebbistatin (pAB) treated cells. 4, 3, 2, 1 asterisk symbols denote P values P < 0.0001, P < 0.001, P < 0.01, and P < 0.05, respectively. The exact P values are shown in Supplementary Table 1.

The above described effects of SMIFH2 inhibition of stress fiber retraction and actin arc movement in permeabilized cells questioned the inhibitory selectivity of SMIFH2 on formin-dependent actin polymerization. Indeed, the permeabilized cells do not contain G-actin, and the incubation buffer was supplied with the actin filament-stabilizing drug, phalloidin. Thus, processes of either polymerization or depolymerization of actin filaments can hardly occur in this system. Given that the effects of SMIFH2 addition phenocopied the action of para-aminoblebbistatin in this assay raised the question of whether SMIFH2 might also be inhibiting nonmuscle myosin 2 paralogs. To address this question, we examined the effect of SMIFH2 on myosin 2A and other myosins *in vitro*.

### Effects of SMIFH2 on myosins *in vitro*

Two methods are primarily used to assess actomyosin function *in vitro*, the actin-activated ATPase activity and the ability of myosin to propel actin filaments in the gliding actin *in vitro* motility assay. In the absence of actin, myosins have very low basal ATPase rates which are activated 10-1000 fold by the addition of actin (De La Cruz and Ostap, 2009). We also used, in some cases, soluble fragments of myosins, termed heavy meromyosin (HMM) for these assays which are considered excellent models for the behavior of the intact myosin, but do not form filaments which complicates the measure of the ATPase activity *in vitro*. We first investigated the effect of SMIFH2 on the actin-activated MgATPase activity of phosphorylated human nonmuscle myosin 2A. The drug inhibited this activity in a dose-dependent manner with an IC_50_ of approximately 50 µM (Fig. 3A). Nonmuscle myosin 2A requires phosphorylation of the regulatory light subunit in order to be activated by actin. The inhibition of the ATPase could potentially be via inhibition of the myosin ATPase activity itself or by inhibition of myosin light chain kinase, which is used to phosphorylate the regulatory light chain of the myosin. However, we found that SMIFH2 did not inhibit the activity of myosin light chain kinase which was used to phosphorylate this myosin (Supplementary table 2). SMIFH2 also inhibited the actin activated ATPase activity of skeletal muscle myosin 2 with an IC_50_ of about 40 µM (Fig. 3B). SMIFH2 inhibited the basal ATPase activity of this myosin in the absence of actin by 89% at 100 µM concentration (Supplementary Figure 1) demonstrating that the drug is acting on myosin and is not inhibiting activity via a direct effect on actin.

**Figure 3:**
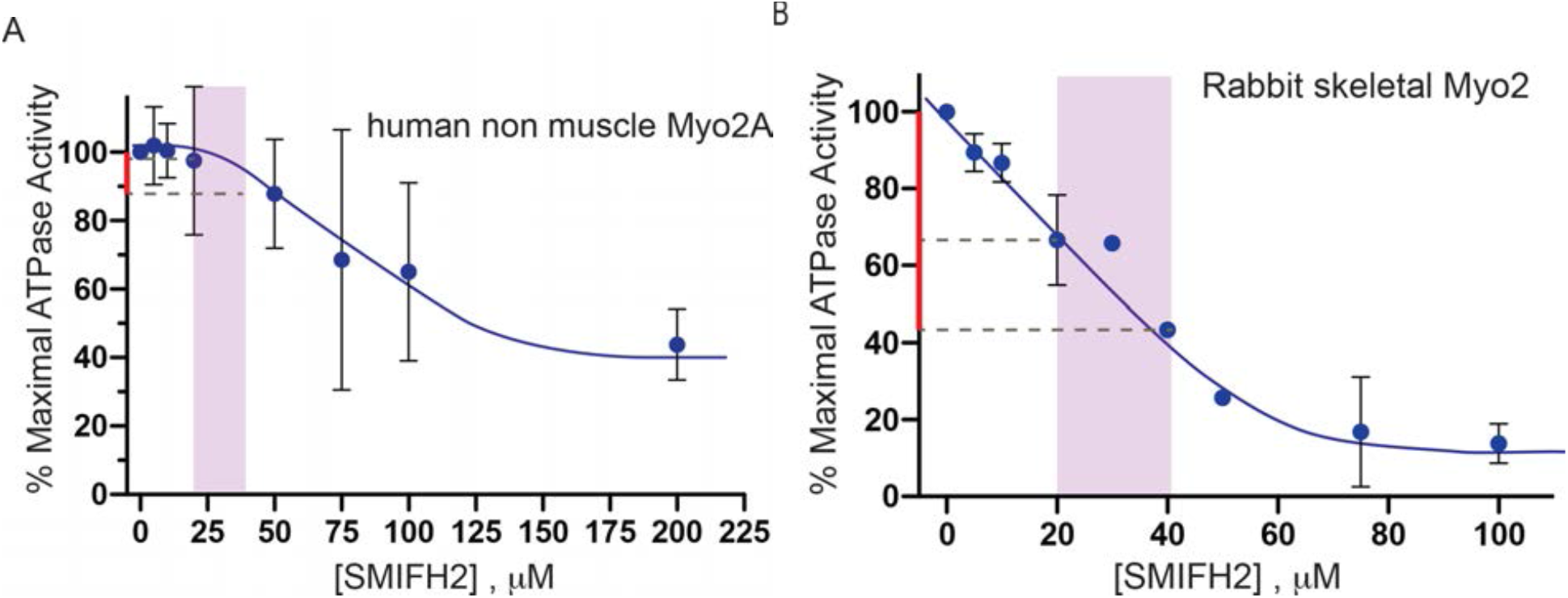
Inhibition of myosin II activities by SMIFH2. A) Inhibition of the actin-activated ATPase of human nonmuscle myosin 2A by SMIFH2; B) inhibition of the actin-activated ATPase of rabbit skeletal muscle myosin 2. The purple area shows to the typical concentration range for SMIFH2 as used in majority of publications. The red lines on the y-axis denote the extent of inhibition approachable at the SMIFH2 concentrations not exceeding these values.

We next examined the effect of SMIFH2 on the ability of skeletal muscle myosin to translocate actin in the gliding actin *in vitro* motility assay. Here skeletal muscle myosin 2 HMM was bound to a nitrocellulose-coated coverslip and its ability to translocate rhodamine-phalloidin labeled actin filaments was observed (Table 1). In the absence of SMIFH2 more than 90% of the actin filaments were motile and moved with a velocity of 5.6 ± 0.6 μm·s^−1^. This activity was completely abolished at an SMIFH2 concentration of 150 µM and we were not able to reverse the inhibition by extensive washout with motility buffer containing no SMIFH2. Interestingly, in these conditions there were immobile actin filaments tethered to the surface, but fewer filaments were present than was observed in chambers that contained no or lower concentrations of SMIFH2. At intermediate SMIFH2 concentrations the average velocity was reduced (Table 1), but the number of motile filaments still remained high. A higher drug concentration was required to bring about 50% of the gliding velocity than was required for similar inhibition of the actin-activated ATPase. This was also observed for blebbistatin inhibition of actin gliding (Limouze et al., 2004; Sakamoto et al., 2005). This fact, coupled with the observation that fewer actin filaments bound to the surface was observed at saturating SMIFH2 concentration suggests that the drug blocks the kinetic cycle of the myosin in a weakly bound state, similar to what was observed for blebbistatin inhibition of myosin 2 isoforms (Kovács et al., 2004; Ramamurthy et al., 2004). Thus, the differences in the concentration of SMIFH2 required for inhibition of the ATPase activity and of the *in vitro* translocation of actin can be explained. The level of inhibition of the ATPase activity is a numerical average of the number of myosins with bound SMIFH2 (fully inhibited) and the ones that do not have drug bound (maximally activated). When 50% of the myosins have bound SMIFH2, the observed actin activated ATPase activity is half maximal. In contrast, in the gliding actin *in vitro* motility assay, the rate of actin filament sliding is not strongly dependent on the number of myosin molecules contributing to movement (the myosin surface density). Therefore, when 50% of the myosins have bound SMIFH2 there are still sufficient active myosins on the surface to propel the actin filaments at full velocity if the inhibited molecules were not able to interact with myosins. However, if SMIFH2 binding blocks the kinetic cycle of myosin to create molecules that can weakly, but not productively, bind to actin, these weakly-bound myosins exert a small frictional drag on the actin filament that will slightly inhibit its velocity (Table 1). This, along with the weakly tethered actin filaments observed on the surface at saturating concentrations of SMIFH2 suggest that this drug, similar to the kinetic action of blebbistatin, likely blocks phosphate release from the acto•myosin•ADP•Pi complex which can only weakly bind to actin and cannot complete the powerstroke.

**Table 1.** Inhibition of skeletal muscle myosin 2 propelled actin filament movement by SMIFH2 in the gliding actin *in vitro* motility assay. Coverslip surfaces were coated with rabbit skeletal muscle myosin 2 HMM (0.2 mg/ml). Motility of rhodamine-phalloidin labeled actin filaments were observed in the absence and presence of the indicated concentrations of SMIFH2. At 150 and 200 μM SMIFH2 no movement of actin filaments was observed and immobile actin filaments were tethered to the surface. Following observation of total inhibition of actin filaments in the presence of 150 μM, the flow cell was extensively washed with 20 volumes motility buffer and then rhodamine-phalloidin actin filaments were re-imaged. There was no movement.

| [SMIFH2], $\mu\text{M}$ | Velocity, $\mu\text{m/s}$ | S.D. , $\mu\text{m/s}$ |
| --- | --- | --- |
| 0 | 5.6 | 0.9 |
| 50 | 4.1 | 0.9 |
| 100 | 3.8 | 0.6 |
| 150 | No movement |  |
| 200 | No movement |  |
| 150 followed by washout | No movement |  |

There are 39 myosin genes from twelve classes in the human genome (Berg et al., 2001). Many, but not all of these myosins are present in other metazoan species including *Drosophila* (Yamashita et al., 2000). While all of these myosins contain a conserved motor domain, the tail portions are very diverse allowing the myosins to perform a plethora of cellular functions. Blebbistatin was shown to be specific for class II myosins (Limouze et al., 2004). To determine whether SMIFH2 inhibits other myosin classes, we assayed its effect on *Drosophila* myosin 5, *Drosophila* myosin 7a and bovine myosin 10. SMIFH2 inhibited the ATPase activity of each of these myosins with various degrees of potency. It inhibited *Drosophila* myosin 7a with an IC_50_ of about 40 µM (Fig. 4A) and myosin 10 with an IC_50_ of about 15 µM (Fig. 4B). Interestingly, SMIFH2 inhibited the activity of *Drosophila* myosin 5 even more potently with an IC_50_ of about 2 µM (Fig. 4C). Thus, SMIFH2 inhibits *Drosophila* myosin 5 even more potently than it does formins *in vitro*. The assays with myosin 10 are particularly interesting since this myosin plays a role in the initiation and formation of filopodia in mammalian cells (Kerber and Cheney, 2011), a process which also involves formin action. The sensitivity of myosin 10 to SMIFH2 questions the use of this drug in the studies of filopodia formation.

**Figure 4.**
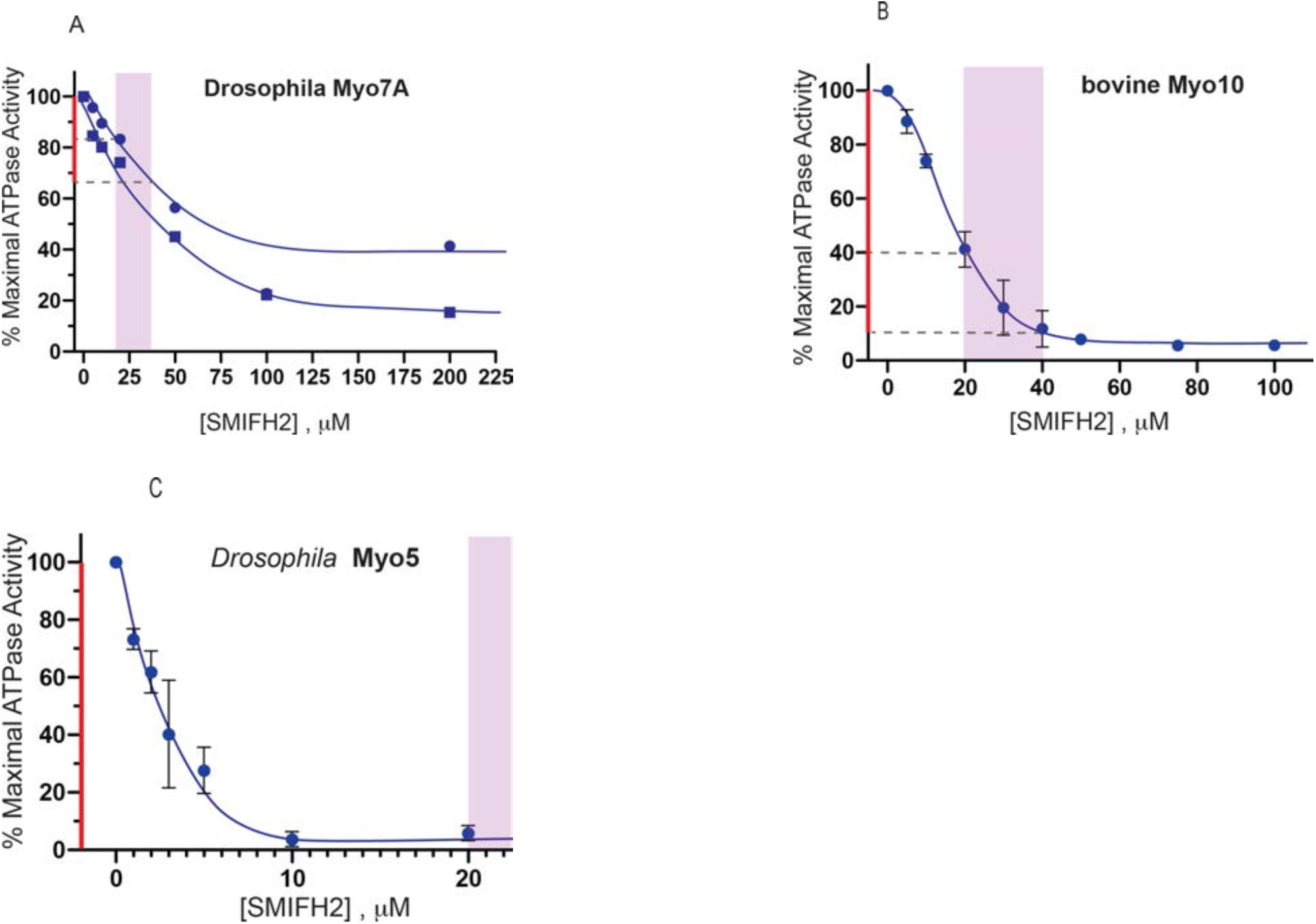
Inhibition of non-conventional myosins by SMIFH2. A) Inhibition of the actin-activated ATPase activity of *Drosophila* myosin 7a; B) inhibition of the ATPase activity of bovine myosin 10; C) inhibition of the ATPase activity of *Drosophila* myosin 5. The purple areas and red lines on y-axis denote the typical concentration range for SMIFH2 and the extent of inhibition, respectively, as explained in the legend to Figure 3.

## Discussion

The actin cytoskeleton of cells consists of many distinct higher order arrays of actin filaments such as the branching network found in the lamellipodia, transverse arc bundles, ventral and dorsal stress fibers and filopodia (Blanchoin et al., 2014; Svitkina, 2018). Some cells contain specialized actin arrays such as microvilli in intestinal enterocytes and stereocilia in the hair cells of the ear (Pelaseyed and Bretscher, 2018). These arrays are formed by actin nucleators such as the Arp2/3 complex, formins and several others with the assistance of a variety of actin associated-proteins (Courtemanche, 2018; Merino et al., 2020; Rottner et al., 2017; Siton-Mendelson and Bernheim-Groswasser, 2017; Swaney and Li, 2016). SMIFH2 was discovered in a small molecule screen for compounds inhibiting formin-driven actin polymerization *in vitro* but not affecting polymerization of pure actin (Rizvi et al., 2009). It has been widely used in the literature to determine the involvement of formins in many cellular and developmental processes and to investigate the immediate effects of formin inhibition. SMIFH2 has been used at concentrations ranging from 5-100 µM and for incubation times of less than 1 hr to more than 24 hrs.

We have found that moderate concentration of SMIFH2 (30μM) very rapidly reduced the traction forces exerted by fibroblasts on their substrates. Further, using cells confined to micropatterned circular adhesive islands, we demonstrated that SMIFH2 inhibits the retrograde flow of myosin filaments in the course of centripetal movement of contracting transverse arcs. These findings by themselves were not alarming since the dynamics of the actin polymerization /depolymerization and interaction of myosins with actin are thought to be tightly linked (Agarwal and Zaidel-Bar, 2019; Zimmermann et al., 2015). However, the fact that SMIFH2 inhibited the ATP-dependent movement of transverse arcs and retraction of ventral stress fibers in permeabilized cells in a manner resembling the blebbistatin inhibition of these processes was unexpected. These permeabilized cells do not contain G-actin, and their actin cytoskeleton is stabilized with phalloidin, which makes the processes of actin polymerization-depolymerization hardly possible. This led us to investigate the possibility that SMIFH2 might inhibit nonmuscle myosin 2A.

Our results show that SMIFH2 inhibits the actin activated ATPase activity of nonmuscle myosin 2A. This was a direct effect on the activity of the myosin since SMIFH2 did not inhibit phosphorylation of myosin 2A light chain by MLCK which is required for nonmuscle myosin 2A activation. It also inhibited both the actin dependent and the basal activity (in the absence of actin) of skeletal muscle myosin 2. The gliding of actin filament by myosin 2A *in vitro* was also suppressed by SMIFH2. Thus, direct inhibition of myosin 2A by SMIFH2 could be involved in the effects of SMIFH2 observed in cells. In our experiments, the concentrations of inhibitor sufficient to stop the myosin 2A filaments flow in living and even permeabilized cells were, however, still lower than the concentrations needed for substantial inhibition of myosin 2A function *in vitro*. In addition, we have recently shown that SMIFH2 can efficiently detach actin filaments from mDia1 formin immobilized on the surface (Alieva et al., 2019). This suggest that SMIFH2 effect on the myosin 2A filaments flow could be at least partially explained by disruption of the actin network connectivity. This possibility deserves further investigation. Nevertheless, nonmuscle myosin 2 paralogs (2A, 2B and 2C) are involved in many cellular processes, which are also thought to involve formins, such as platelets formation (Pal et al., 2020; Pan et al., 2014; Zhang et al., 2012b), assembly of the cytokinetic contractile ring (Pollard and O’Shaughnessy, 2019; Taneja et al., 2020) and maintenance of stress fiber integrity (Hu et al., 2017; Oakes et al., 2012). Thus, the conclusions about formin function based solely on experiments with SMIFH2 should be carefully revisited.

Even more surprising were the findings that SMIFH2 affects some other types of myosins stronger than myosin 2A and in some cases stronger than it affects formins. We have demonstrated that SMIFH2 inhibits myosins from all classes that were tested including rabbit skeletal muscle myosin, *Drosophila* myosins 5 and 7a and mammalian myosin 10. The IC_50_ values for inhibition of myosin 10 were similar to that for the inhibition of formins. *Drosophila* myosin 5 was inhibited with an IC_50_ of about 2 µM, which is more potent than the published values for a variety of formins.

The actions of formins and myosin are intimately linked in cells since most of the actin arrays built or influenced by formins interact with myosins. For example the cytokinetic ring formation depends on formins (Pollard and O’Shaughnessy, 2019), but nonmuscle myosin 2 paralogs are essential for its function (Taneja et al., 2020; Yamamoto et al., 2019). Filopodia elongation is proven to be formin-dependent in formin knockdown and overexpression experiments (Mellor, 2010; Schaks et al., 2019), but many studies demonstrate that myosin 10 also plays an important role. Myosin 10 was found in the patches at the tips of filopodia in mammalian cells and its knockdown in a variety of cells is associated with filopodia suppression (Arjonen et al., 2011; Kerber and Cheney, 2011). In addition, nonmuscle myosin 2A was shown to play a role in the stabilization of filopodia adhesion (Alieva et al., 2019). In our recent experiments, effect of SMIFH2 on filopodia involves disintegration of myosin 10 patches at filopodia tips and myosin 2A-dependent centripetal movement of residual myosin 10 puncta along filopodia (Alieva et al., 2019). Thus, SMIFH2 treatment did not inhibit myosin 2A activity in these experiments, but its effect on myosin 10 cannot be excluded. Thus, experiments with SMIFH2 alone do not permit to dissect functions of formins and myosins in filopodia.

Some mutations in DIAPH1 correlate with hearing loss and there are some evidences of formin involvement in formation of stereocilia in specialized inner ear cells (Neuhaus et al., 2017; Ueyama et al., 2016). At the same time, the formation and function of these same structures are dependent on a variety of myosin isoforms, including myosin 1C (Stauffer et al., 2005), myosins 3a and b (Lelli et al., 2016), myosin 6 (Hertzano et al., 2008; Seiler et al., 2004), myosins 7a (Morgan et al., 2016; Yu et al., 2017), and myosin 15 (Anderson et al., 2000; Friedman et al., 1999). Formation and maintenance of integrity of actin bundles (radial/dorsal fibers, transverse arcs and ventral stress fibers) depend on formins (Hotulainen and Lappalainen, 2006; Oakes et al., 2012; Schulze et al., 2014), but these structures depend also on myosin 2 driven force generation, and nonmuscle myosin 2 paralogs are components of some of these structures (Beach et al., 2014; Hu et al., 2017; Kuragano et al., 2018; Shutova et al., 2017; Vicente-Manzanares et al., 2009). Myosin 2 and 7a has also conserved function in cell adhesions (Küssel-Andermann et al., 2000; Titus, 2005; Velichkova et al., 2002; Vicente-Manzanares et al., 2009), a process in which formins are also participating (Grikscheit and Grosse, 2016; Romero et al., 2020). Thus, it would be difficult to interpret studies of formin functions using a compound that inhibits both formins and myosins.

An unsolved question is whether there is some structural similarity between myosins and formins which could explain dual specificity of SMIFH2. Such similarity, however, does not necessary exists, since the highly electrophilic nature of SMIFH2 makes this chemical very promiscuous in its interactions with different proteins, as was recognized in screening-based studies (Baell, 2010).

In summary, our study demonstrates that SMIFH2 can no longer be considered as a specific inhibitor of formins in the studies related to cell motility and actomyosin cytoskeleton organization. The conclusions based on using of SMIFH2 in such studies should be carefully reconsidered and possibly reinterpreted. The development of novel more specific inhibitors suitable for instant suppression of formin functions in cells is becoming an important and timely task for future studies.

## Materials and Methods

### Cell culture and transfection

The immortalized rat embryo fibroblasts (REF52 cells) cell line (Matsumura et al., 1983) and Human foreskin fibroblasts (HFFs) (American Type Culture Collection, Manassas, VA, USA; catalogue no. SCRC-1041) were cultured in Dulbecco’s modified Eagle’s medium (DMEM; Invitrogen, 11965092) supplemented with 10% heat-inactivated fetal bovine serum (FBS; Invitrogen, 10082147) and 1% penicillin/streptomycin (Invitrogen, 15070063) at 37°C and 5% CO_2_. Both cell lines were regularly tested for mycoplasma contamination by MycoAlert PLUS Mycoplasma Detection Kit (Lonza, LT07-703). REF52 cells were transiently transfected with mEos3.2-Actin expression vector (Michael W. Davidson group collection, The Florida State University, Tallahassee, FL, USA, kindly provided by Dr. P. Kanchanawong, MBI) by jetPRIME transfection reagent (Polyplus transfection, 114-15) in accordance with the manufacturer’s protocols. HFF cells were transfected with myosin regulatory light chain (RLC)-GFP expression vector (Kengyel et al., 2010) (a gift from W. A. Wolf and R. L. Chisholm, Northwestern University, Chicago, IL, USA) using electroporation (Neon transfection system, Life Technologies) following the manufacturer’s instructions.

### Traction force microscopy

The traction force microscopy with embedded beads is performed as described previously (Rafiq et al., 2019). Briefly, a soft polydimethylsiloxane CY 52-276A and CY 52-276B (Dow Corning, 0008602722) were mixed with the ratio 1:1 and the Sylgard 184 crosslinker was used to tune the stiffness of the gel for proper force measurement of cells (∼95 kPa). The mixture was spin-coated onto a clean coverslip to achieve the thickness of ∼7μm and cured for 1 h at 80 °C. The surface of the gel was silanized with (3-aminopropyl) triethoxysilane for 2 h, followed by incubation of 0.04μm carboxylate-modified dark red (660/680) beads (Thermo Fisher Scientific, 1871942) at 1 × 10^6^ beads/ml in a solution of 0.1 M sodium bicarbonate for 30 min. Before seeding the cells, the coverslips with beads were further incubated for 30 min with 10 μg/ml fibronectin also dissolved in 0.1 M sodium bicarbonate. The traction forces were calculated from bead displacement field visualized by live cell imaging as described in Tseng et al (2012) using the online ImageJ plugin (https://sites.google.com/site/qingzongtseng/tfm for plugin software details). The computation algorithm by Sabass et al (2008) was used. The distribution of traction force magnitude was presented as a heat map (Fig. 1A). The mean magnitude value was calculated for each cell.

### Cytoskeleton contractility assays

For HFF cells, circular adhesive islands of fibronectin were fabricated by stencil patterning described previously (Jalal et al., 2019). GFP-MLC expressing HFF cells were then seeded at density of 5 × 10^4^ cells/ml on the hydrophobic uncoated 35mm μ-dish (ibidi,81151) with fibronectin micro-patterns and incubated 3-8 hours prior to the experiment. For stress fiber imaging, REF52 cells were transfected with mEos3.2-Actin expression plasmid and seeded onto a 35mm glass bottom dish (Iwaki, 3930-035) 24 hours prior to the assay.

The protocol of the cell permeabilization and cytoskeleton contractility assay was described previously (Tee and Bershadsky, 2016; Tee et al., 2015). Briefly, cells were permeabilized with extraction buffer A (50mM imidazole (pH 6.8), 50mM KCl, 0.5 mM MgCl_2_, 0.1mM EDTA, 1mM EGTA, 1mM 2-Mercaptoethanol, 250nM phalloidin (Thermo fisher, P34572) and 2μg/mL protease inhibitor cocktail (Sigma, P8340)) supplemented with 0.1% Triton-X100 and 4% PEG MW35000 for 10 min at room temperature, then washed three times with extraction buffer A. The cytoskeleton contractility assay was carried out at 37°C with buffer A supplemented with 2mM ATP with or without the appropriate drugs. SMIFH2 (Sigma, S4826) and para-aminoblebbistatin (pAB, Optopharma, DR-Am-89) were used. All drugs were remained in the buffer during the entire period of observation.

### Live cell imaging and confocal microscopy

Super-resolution SIM imaging was performed using W1-spinning-disc confocal unit coupled with the live super-resolution (SR) module (spinning disk based structured illumination super resolution (York et al., 2013), GatacaSystems), mounted on Eclipse microscope with Perfect Focus System, supplemented with the objective Plan Apo 100x oil NA1.45 or 60x 1.20 NA CFI Plan Apo Lambda water immersion (Nikon) and scientific complementary metal–oxide–semiconductor (sCMOS) camera Prime95B (Photometrics). Laser lines wavelength 488, 561 and 647nm were used. For HFF cells, time-lapse images at 2 min intervals of Z-stacks with step-size 0.35 μm were acquired. For REF52 cells, time-lapse images at 5 min intervals at the basal plane of the cells were acquired for 30min.

### Image analysis

Particle Image Velocimetry (PIV) analysis was used for measuring average instantaneous speed of the GFP-Myosin light chain labeled myosin filaments. First, the maximum projection of time lapse of GFP channel was segmented using in-house MATLAB code to get the mask of region-of-interest (ROI). After segmentation, PIV analysis was performed using MatPIV 1.6.1 (https://www.mn.uio.no/math/english/people/aca/jks/matpiv/). Single-pass PIV with window size of 32×32 pixels and 50% overlapping was applied. The average instantaneous speed for the first two frames within the ROI was computed. To quantify the retraction speed of stress fiber ends, we manually selected all ends labeled by unconverted mEOS3.2-actin and arranged them into kymographs for every cell. The retraction speed for each end during imaging was calculated manually using the kymograph. The average retraction speed of stress fibers was calculated for each cell and plotted in the graph.

### Statistical analyses

Plotting and statistical analysis were done using GraphPad Prism 7 (GraphPad Software). The significance of the differences (P value) was calculated using two-tailed unpaired Student’s t-test. Bar graphs and scatter plots show mean±s.d. for the respective groups of data.

### Preparation of proteins

A heavy meromyosin (HMM)-like fragment of human nonmuscle myosin 2A was prepared by expression in Sf9 cells as described (Kengyel et al., 2010). The cells were co-infected with a virus driving the expression of the truncated myosin heavy chain as well as one that drove the expression of RLC and ELC. The myosin was phosphorylated with smooth muscle MLCK prior to use (Nagy et al., 2013). Full length *Drosophila* myosin 5 (Lu et al., 2020), *Drosophila* myosin 7a (Yang et al., 2009) and a forced dimeric HMM-like fragment of bovine myosin 10 (Takagi et al., 2014) were also produced in Sf9 cells. *Drosophila* myosin 5 was co-expressed with *Drosophila* ELC and calmodulin whereas the myosin 10 was co-expressed with calmodulin. Skeletal muscle HMM was produced by chymotrypic digestion of full length rabbit fast skeletal muscle myosin (Margossian and Lowey, 1982). Rabbit skeletal muscle actin was prepared as previously described (Lehrer and Kerwar, 1972).

### Actin-activated ATPase assays

ATPase assays were carried out using an NADH-linked assay in a Cary 50 spectrophotometer as previously described (Heissler et al., 2015). SMIFH2 was first prepared as an 50 mM solution in DMSO and was diluted in DMSO as required. The final amount of DMSO added to the samples was 2%. The assay conditions were 10 µM actin, 50 mM KCl, 10 mM MOPS, 2 mM MgCl_2_, 0.1 mM EGTA, 1 mM ATP, 10 mM MOPS (pH 7.0) at 25°C. The buffer 40 units/ml l-lactic dehydrogenase, 200 units/ml pyruvate kinase, 200 μm NADH, and 1 mm phosphoenolpyruvate. The absorbance was monitored at 340 nm. Nonmuscle myosin 2A was first phosphorylated by incubation in 0.3 M KCl, 4 mM MgCl_2_, 0.2 mM CaCl_2_, 0.1 mM EGTA, 0.1 mM ATP, 1 μM calmodulin, 2 nM myosin light chain kinase for 10 min at room temperature.

### Gliding actin in vitro motility assay

The gliding actin in vitro motility assay was conducted at 30 °C in 50 mM KCl, 4 mM MgCl_2_, 0.1 mM EGTA, 0.5% methylcellulose, 1 mM ATP with an oxygen savaging system consisting of 2.5 μg/ml glucose oxidase, 45 μg/ml catalase, 2.5 mg/ml glucose, and 50 mm DTT (Sellers et al., 1993). The rate of movement of actin filaments was determined as described previously (Homsher et al., 1992).

### HPLC-Mass spectrometry for Protein Phosphorylation

Phosphorylation of NM2A was initiated by the addition of ATP. Samples were taken at different time points and diluted with 305 acetonitrile, 0.25 TFA to stop the reaction. Proteins were injected into a reverse phase HPLC (Agilent 1100 series HPLC, Agilent Technologies) with a Zorbax 300SB-C18 (2.1 x 50mm, 3.5mm, Agilent Technologies) and introduced into the mass spectrometer as described (Apffel et al., 1995; Taggart et al., 2000). Positive ion Electrospray Ionization (ESI) mass spectra for intact protein were obtained with an Agilent 6224 mass spectrometer equipped with an ESI interface and a time-of-flight (TOF) mass detector (Agilent Technologies). Mass spectra were analyzed and de-convoluted using a software, MassHunter version B.06.00 (Agilent Technologies).

## Acknowledgements

We are grateful to Dr. Yee Han Tee (MBI, Singapore) for providing detailed protocol for experiments with permeabilized cells and to Ms. Hui Ting Ong (MBI, Singapore) for developing the software for PIV data analysis. We acknowledge the NHLBI Biochemistry Core for performing mass spec analysis of regulatory light chain phosphorylation. A.D.B. acknowledges the support from the Singapore Ministry of Education Academic Research Fund Tier 2 (MOE Grant No: MOE2018-T2-2-138), the National Research Foundation, Prime Minister’s Office, Singapore, and the Ministry of Education under the Research Centres of Excellence programme through the Mechanobiology Institute, Singapore (ref no. R-714-006-006-271), and Singapore Ministry of Education Academic Research Fund Tier 3 MOE grant no. MOE2016-T3-1-002. JRS is funded by HL001786.

## Figures

**Supplementary Table 1.**

|  | SMIFH2 | pAB |
| --- | --- | --- |
| Non-treated | <0.0001 | <0.0001 |

(2) P values in Figure 1D
| | 30 $\mu$ M SMIFH2 | 50 $\mu$ M SMIFH2 | 100 $\mu$ M SMIFH2 | 200 $\mu$ M SMIFH2 | 100 $\mu$ M pAB |
| --- | --- | --- | --- | --- | --- |
| Non-treated | 0.0030 | <0.0001 | <0.0001 | <0.0001 | <0.0001 |
| 100 $\mu$ M pAB | <0.0001 | <0.0001 | 0.0003 | 0.0107 | |

(3) P values in Figure 2G
| | 12.5 $\mu$ M SMIFH2 | 25 $\mu$ M SMIFH2 | 50 $\mu$ M SMIFH2 |
| --- | --- | --- | --- |
| Non-treated | 0.0002 | <0.0001 | <0.0001 |

| | 12.5 $\mu$ M SMIFH2 | 25 $\mu$ M SMIFH2 | 50 $\mu$ M SMIFH2 | 100 $\mu$ M SMIFH2 | 100 $\mu$ M SMIFH2 | 100 $\mu$ M pAB |
| --- | --- | --- | --- | --- | --- | --- |
| Non-treated | 0.0002 | <0.0001 | <0.0001 | <0.0001 | <0.0001 | <0.0001 |
| 100 $\mu$ M pAB | <0.0001 | <0.0001 | 0.2153 | 0.0009 | 0.7841 | |

**Supplementary Table 2.** SMIFH2 has no effect on myosin light chain kinase. Nonmuscle myosin 2A was incubated with myosin light chain kinase in the presence or absence of 100 μM SMIFH2 for the indicated times. The extent of phosphorylation was quantified by mass spectrometry (Apffel et al., 1995; Taggart et al., 2000).

| | 0 μM SMIFH2 | 100 μM SMIFH2 |
| --- | --- | --- |
| Time, min. | % phosphorylated | % phosphorylated |
| 0.5 | 50.8 | 55.4 |
| 1.5 | 71.2 | 70.4 |
| 4.5 | 82.1 | 81.2 |

